# Discovery of a human testis-specific protein complex TEX101-DPEP3 and selection of its disrupting antibodies

**DOI:** 10.1101/315713

**Authors:** Christina Schiza, Dimitrios Korbakis, Efstratia Panteleli, Keith Jarvi, Andrei P. Drabovich, Eleftherios P. Diamandis

**Affiliations:** Department of Laboratory Medicine and Pathobiology, University of Toronto, Toronto, Canada; Department of Pathology and Laboratory Medicine, Mount Sinai Hospital, Toronto, Canada; Lunenfeld-Tanenbaum Research Institute, Mount Sinai Hospital, Toronto, Canada; Department of Clinical Biochemistry, University Health Network, Toronto, Canada; Department of Surgery, Division of Urology, Mount Sinai Hospital, Toronto, Canada

**Keywords:** TEX101, testis-expressed sequence 101 protein, DPEP3, dipeptidase 3, affinity purification-mass spectrometry, interactome, protein-protein interactions, testis-specific proteins, spermatozoa, disrupting antibodies

## Abstract

TEX101 is a testis-specific protein expressed exclusively in male germ cells and is a validated biomarker of male infertility. Studies in mice suggest that TEX101 is a cell-surface chaperone which regulates, through protein-protein interactions, the maturation of proteins involved in spermatozoa transit and oocyte binding. Male TEX101-null mice are sterile. Here, we identified by co-immunoprecipitation-mass spectrometry the interactome of human TEX101 in testicular tissues and spermatozoa. The testis-specific cell-surface dipeptidase 3 (DPEP3) emerged as the top hit. We further validated the TEX101-DPEP3 complex by using hybrid immunoassays. Combinations of antibodies recognizing different epitopes of TEX101 and DPEP3 facilitated development of a simple immunoassay to screen for disruptors of TEX101-DPEP3 complex. As a proof-of-a-concept, we demonstrated that anti-TEX101 antibody *T4* disrupted the native TEX101-DPEP3 complex. Disrupting antibodies may be used to study the human TEX101-DPEP3 complex, and to develop modulators for male fertility.

**Non-standard abbreviations:**

TEX101: Testis-expressed sequence 101 protein
DPEP3: Dipeptidase 3
AC-MS: Affinity capture-mass spectrometry
co-IP-MS: Coimmunoprecipitation-mass spectrometry
GPI: Glycosylphosphatidylinositol
LFQ: Label-free quantification
mAb: Monoclonal antibody
NHS: N-hydroxysuccinimide
PRM: Parallel reaction monitoring
PTM: Posttranslational modification
SP: Seminal plasma
SRM: Selected reaction monitoring
FDR: false detection rate

## INTRODUCTION

Spermatogenesis is a highly organized process involving coordinated cell cycle progression, differentiation of spermatogonial stem cells and their transformation into mature spermatozoa. With no cell culture models of human germ cells available as yet, the molecular biology of spermatogenesis remains one of the least studied developmental processes in humans.

Numerous animal studies emphasized the importance of protein-protein interactions (PPIs) in the production of fertile spermatozoa. In fact, the necessity to silence transcription and translation at the late stages of spermatogenesis resulted in the evolution of epididymis, in which spermatozoa are activated by epidydimis-secreted proteins through numerous proteolytic cascades and PPIs. Null mice models of selected testis-specific genes presented with male infertility phenotypes, presumably through disrupted PPIs and improper processing of proteins during spermatogenesis and sperm maturation (1-7). Early studies discovered the essential role of numerous cell surface proteins for sperm-oocyte interaction and fusion (8). Some of the most critical factors included metalloprotease-disintegrin ADAM2 (9), the cell adhesion tetraspanin CD9 (10) and the sperm-egg fusion protein IZUMO1 (11). The recent discovery of the cell surface recognition complex of IZUMO1 protein and the sperm-egg fusion protein JUNO provided detailed insights into gamete recognition and sperm-oocyte fusion (12, 13). Of 1,035 highly testis-enriched proteins in the human proteome (14), nearly 160 proteins are membrane-bound and could be involved in spermatogenesis, remodelling of spermatozoa cell surface, sperm transit and sperm-oocyte interaction. The identification of the exact roles of PPIs during maturation of male and female germ cells continues.

Proteomics and mass spectrometry emerged as the techniques of choice to discover PPIs and to elucidate the molecular functions of proteins (15-17). Affinity purification or co-immunoprecipitation (co-IP) approaches followed by mass spectrometry identified numerous direct and indirect PPIs under native physiological conditions (18-20). Advances in sensitivity and throughput of mass spectrometry facilitated mapping of interactomes of bacteria (21), yeast (22, 23), insects (24) and human cells (25). High resolution mass spectrometry empowered by label-free quantification enabled identification of high-confidence PPIs after a single step of affinity purification (26).

In this study, we focused on the testis-specific protein TEX101, which we previously discovered and validated as a biomarker of male infertility (27-31). TEX101 protein is exclusively expressed on the surface of testicular germ cells (32) and was suggested to be a cell-surface chaperone involved in trafficking and maturation of numerous cell surface proteins essential for fertilization in mice (7, 33). With four TEX101-regulated proteins (ADAM3-6) previously discovered in mice (7, 34), three correspond to pseudogenes in humans, while ADAM4 gene is not present in the human genome. While human TEX101 is an outstanding biomarker of male infertility (28), its functional role in human reproduction is not known. In this work, we established a quantitative co-IP-MS approach to discover the human TEX101 interactome. Taking into account the degradation of testis-specific ADAM proteins in TEX101-null mice and subsequent sterility of male mice (7), we hypothesized that disruptors of TEX101 PPIs could emerge as modulators of male fertility and non-hormonal male contraceptives.

## EXPERIMENTAL PROCEDURES

### Expression and purification of recombinant proteins TEX101 and DPEP3

Human recombinant proteins TEX101 and DPEP3 were produced in an Expi293F transient expression system according to manufacturer’s recommendations (Invitrogen). Briefly, DNA coding for the mature forms of TEX101 and DPEP3 (aa 26-222 and 36-463, respectively) were cloned into a pcDNA3.4 plasmid for mammalian protein expression (GeneArt^TM^ Gene Synthesis, Invitrogen). Expi293F cells were grown in suspension and cell cultures containing secreted TEX101 and DPEP3 proteins were collected 72 and 96 hours post-transfection, respectively. Recombinant protein production was assessed by Western blot analysis with rabbit polyclonal antibodies, anti-TEX101 HPA041915 (Sigma-Aldrich) and anti-DPEP3 HPA058607 (Sigma-Aldrich). Purification of recombinant TEX101 and DPEP3 from culture supernatants was performed with an automated AKTA FPLC system on a pre-equilibrated 5-mL anion-exchange HiTrap Mono Q^TM^ Sepharose high performance column (GE Healthcare). Culture supernatants were diluted in 50 mM Tris (buffer A), pH 9.0 for TEX101, and pH 9.5 for DPEP3, and following binding and washing, proteins were eluted in 4-mL fractions with a linear gradient of 50 mM Tris, 1 M NaCl (buffer B), pH 9.0 (TEX101), and pH 9.5 (DPEP3). The concentration of TEX101 and DPEP3 in fractions was measured by TEX101 ELISA (35), and an SRM assay, respectively. Purity and molecular mass of recombinant proteins were determined by SDS-PAGE stained with Coomassie Blue. Gel bands were excised, subjected to in-gel digestion and analyzed by LC-MS/MS in a Q Exactive^TM^ Plus Hybrid Quadrupole-Orbitrap^TM^ Mass Spectrometer (Thermo Scientific). For protein identification, the LC-MS/MS raw files were analyzed using the MaxQuant software (version 1.5.2.8) with the human UniProtKB/Swiss-Prot database (HUMAN5640_sProt-012016).

### Monoclonal antibody production against human TEX101 and DPEP3

All animal research was approved by the TCP Animal Care Committee (Animal Use Protocol #14-04-0119aH). Monoclonal antibody (mAb) production was performed as previously described (35). Female BALB/c mice were inoculated with purified recombinant proteins, TEX101 or DPEP3, and three booster injections were performed at 3-week intervals. After successful fusion of spleen cells with NSO murine myeloma cells, cell culture supernatants were tested for IgG and IgM antibody secretion using an immunoassay protocol described elsewhere (35). In-house developed mouse monoclonal antibodies included: anti-TEX101 antibodies 34ED556 (antibody *T1*), 34ED233 (antibody *T*5) and 34ED470 (antibody *T6*) recognizing epitope A; 34ED229 (antibody *T2*), 34ED629 (antibody *T3*) and 34ED604 (antibody *T4*) recognizing epitope B. Anti-DPEP3 monoclonal antibodies included: 40ED139 (antibody *D1*) recognizing epitope A and 41ED68 (antibody *D2*) recognizing epitope B.

### Immunocapture-SRM screening for clones producing antibodies against native TEX101 and DPEP3 proteins

Screening for clones producing antibodies against native TEX101 and DPEP3 was performed according to our established protocol (35). A commercial mouse polyclonal anti-TEX101 antibody (ab69522; Abcam, Cambridge, MA) was used as a positive control for anti-TEX101 antibody secreting clone screening. Immunocapture-SRM was used for the screening of hybridoma culture supernatants for antibodies against native TEX101 in testicular tissue lysate pool. Prior to MS analysis, a mix of 50 mM ammonium bicarbonate (ABC), 5 mM dithiothreitol (DTT) (Sigma-Aldrich), 100 fmoles of heavy isotope-labeled TEX101 proteotypic peptide AGTETAILATK*-JPTtag with a trypsin-cleavable tag, and 0.05% RapiGest SF was added to each well. Following protein reduction and alkylation, samples were digested overnight at 37°C with the addition of proteomics-grade porcine trypsin (Sigma-Aldrich, #T6567). Trypsin inactivation and RapiGest cleavage were achieved by adding trifluoroacetic acid (TFA) at final concentration of 1%. A two-step IP-SRM was also used for the screening of hybridoma culture supernatants for mAbs against recombinant human DPEP3, and native DPEP3 protein in SP. Serum of immunized mice was used as a positive control. Heavy isotope-labeled DPEP3 proteotypic peptide SWSEEELQGVLR*-JPTtag (300 fmol) was added on the plate, and samples were prepared for mass spectrometric analysis, as described above. TEX101 or DPEP3 peptides were monitored in a non-scheduled SRM mode during a 30 min LC gradient in TSQ Quantiva^TM^ triple quadrupole mass spectrometer (Thermo Scientific). Raw files for each sample were analyzed with Skyline software (v3.6.0.10493), and relative abundance of TEX101 and DPEP3 were calculated using the ratio of endogenous versus internal standard peptides. Hybridoma cultures, positive for antibody secretion against native TEX101 and DPEP3, were grown and transferred in serum-free media (Invitrogen). Supernatants were harvested and purified using protein G according to the manufacturer’s protocol (GammaBind Plus, GE Healthcare).

### Pairing of anti-TEX101 monoclonal antibodies in a sandwich immunoassay

Pairing of purified anti-TEX101 mAbs (*T2*, *T5*, *T6*, *T1*, *T4* and *T3*) was performed as previously described (35). Seminal plasma pool sample diluted 50-fold in assay diluent was loaded on the plates and assay diluent alone was used as a negative control. After 2 hours of incubation, plates were washed, and biotinylated mouse monoclonal anti-TEX101 antibodies in the assay diluent (250 ng per well) were added and incubated for 1 hour. All mAbs were paired with each other in a sandwich format, generating 36 combinations (6x6). After the addition of streptavidin-conjugated alkaline phosphatase, diflunisal phosphate (DFP) solution in substrate buffer, and lastly, developing solution were added. Time-resolved fluorescence was measured with the Wallac EnVision 2130 Multilabel Reader (Perkin Elmer).

### Testicular tissue, spermatozoa and SP samples

Testicular tissues with active spermatogenesis (confirmed by histological examination) were obtained with informed consent by orchiectomy from men with scrotal pain or testicular masses. Upon removal, testicular tissues were subjected to snap-freezing, and stored in liquid nitrogen. Semen samples were collected from healthy fertile pre-vasectomy patients, they were allowed to liquefy at RT for 1 hour and then aliquoted and centrifuged 3 times at 13,000 g for 15 min at RT. The SP and sperm cells were separated and stored at -80°C. Sample collection was approved by the institutional review boards of Mount Sinai Hospital (testicular tissue; approval #09-0156-E and semen; approval #08-117-E) and University Health Network (semen; #09-0830-AE).

### Preparation of testicular tissue and spermatozoa lysates

Testicular tissue and spermatozoa lysis and solubilization of protein complexes was performed under optimized lysis conditions. Cryogenic tissue lysis was followed by suspension of the frozen nsample powder in the lysis buffer containing 20 mM HEPES (pH 7.5), 100 mM NaCl, 1% w/v 3-[(3-cholamidopropyl)-dimethylammonio]- 1-propanesulfonate (CHAPS), and 1% v/v protease inhibitor cocktail [1:10 (w/v) ratio of tissue to lysis buffer]. Several sperm cell samples were pooled and incubated with lysis buffer. After overnight incubation at 4°C, testicular tissue and spermatozoa lysates were centrifuged at 15,000 g for 10 min at 4°C, and total protein concentration was measured by the bicinchoninic acid assay (BCA). Testicular tissue and sperm cell lysates were stored at -20°C.

### Immobilization of IgG antibodies on N-hydroxysuccinimide (NHS)-activated sepharose beads

Two in-house generated mouse monoclonal anti-TEX101 antibodies (T1 and T2) that recognized different epitopes, and a non-specific mouse IgG (isotype control) were immobilized on NHS-activated Sepharose 4 Fast Flow (GE Healthcare), by using a previously optimized protocol (36). Following antibody coupling, sepharose beads were incubated in blocking buffer, and then they were washed with binding buffer 1x TBS (50 mM Tris, 150 mM NaCl, pH 7.5).

### Co-IP of TEX101 complexes in testicular tissues, spermatozoa and SP

Co-IP of TEX101 complexes from testicular tissue lysate (600 μg of total protein) was performed in triplicates with anti-TEX101 antibodies *T1* or *T2* and non-specific mouse IgG coupled to beads (50 μL). Co-IP of TEX101 complexes from spermatozoa lysate (120 μg total protein) was performed in triplicates with *T1* and non-specific mouse IgG coupled to beads (30 μL). Co-IP of TEX101 complexes from SP (600 μg total protein) was performed in triplicates with *T1* and non-specific mouse IgG coupled to beads (50 μL). Following binding for 2 hours at RT with shaking, all beads were washed with TBS binding buffer (50 mM Tris, 150 mM NaCl, pH 7.5) and 50 mM ammonium bicarbonate, and then were re-suspended in 50 mM ammonium bicarbonate. Proteins were subjected to reduction (DTT; 5 mM final), alkylation (iodoacetamide; 10 mM final), and overnight digestion with trypsin (0.5 μg). Supernatants were collected and remaining beads were incubated again with 30% acetonitrile at RT for 10 min. First and second supernatants were pooled, and trypsin was inactivated by 1% TFA.

### Identification of TEX101 complexes by liquid chromatography – tandem mass spectrometry

Following digestion, peptides were extracted with C18 OMIX tips, and samples were analysed by an EASY-nLC 1000 system (Thermo Fischer Scientific) coupled online to a Q Exactive^TM^ Plus Hybrid Quadrupole-Orbitrap^TM^ Mass Spectrometer (37). Each immunoprecipitation full-process replicate was analyzed with an 18 μL single injection. Peptides in each sample were loaded and separated with a 15 cm C18 analytical column (inner diameter 75 μm, tip diameter 8 μm) using a 60-min LC gradient. A data-dependent mode was utilized to acquire a full MS1 scan from 400 to 1500 m/z in the mass analyzer at resolving power of 70,000, followed by 12 precursor ions data-dependent MS2 scans at 17,500 resolution. Ions with charge states of +1, ≥+4, and unassigned charge states were excluded from MS2 fragmentation.

### Experimental design and statistical rationale for identification of TEX101 complexes by LC-MS/MS

Co-IP of TEX101 complexes was performed in pools of testicular tissue lysates, spermatozoa lysates and SP (one biological replicate for each type of specimen). Three full process replicates were performed independently (from co-IP to trypsin digestion) for each specimen, and each process replicate was analysed by a single LC-MS/MS technical replicate. Non-specific mouse IgG was used as a negative control. For protein identification and data analysis, mass spectra, generated by XCalibur (v. 2.0.6; Thermo Fischer Scientific), were processed with MaxQuant software (version 1.5.2.8). Protein search was performed against the non-redundant Human UniProtKB/Swiss-Prot database (HUMAN5640_sProt-012016). Search parameters included: trypsin enzyme specificity, 2 missed cleavages, minimum peptide length of 8 amino acids, minimum of 1 unique peptide, top 8 MS/MS peaks per 100 Da, peptide mass tolerance of 20 ppm for precursor ion and MS/MS tolerance of 0.5 Da, fixed modification of cysteines by carbamidomethylation and variable modification of methionine oxidation and N-terminal protein acetylation. False-discovery rate (FDR) was set to 1% both at the protein and the peptide levels. Label-free relative quantification of identified proteins was achieved by the MaxLFQ algorithm integrated into MaxQuant (38). The ‘proteinGroups.txt’ file, generated by MaxQuant, was uploaded to Perseus software (version 1.5.5.3) for further statistical analysis. Protein identifications classified as “Only identified by site”, “Reverse”, and “Contaminants” were excluded. LFQ intensities were log2-transformed, and two groups with three replicates each were compared (LFQ-anti-TEX101 and LFQ-mouse IgG). Proteins with less than three valid values in at least one group were filtered out. Missing LFQ values were imputed with values representing a normal distribution to enable statistical analysis. A two-sample *t*-test (Benjamini-Hochberg false-discovery rate-adjusted *p* values) was applied to determine proteins statistically enriched by anti-TEX101 versus non-specific mouse IgG. We performed variance correction (s0) for each comparison, and we applied FDR of 1% for candidate selection. Volcano plots were generated to facilitate data visualization. The list of putative TEX101-interacting proteins was merged with the Human Protein Atlas (v.13) secretome (n=2928) and membrane-bound proteome (n=5463), to select secreted and membrane-bound proteins expressed in testis (14). Expression and localization of each candidate protein was manually assessed using Human Protein Atlas immunohistochemistry data and the NeXtProt database.

### Experimental design and rationale for the verification of TEX101-interacting proteins by targeted MS

To verify TEX101 interactome, we developed and applied a Tier 2 targeted mass spectrometry analysis. Two multiplexed SRM assays combined with co-IP were used to monitor the candidate proteins in testicular tissue and spermatozoa. Targeted SRM assays were developed as previously described (39-44). Our MS and MS/MS identification data (including potential post-translational modifications) was used to select proteotypic peptides. Peptides with 7-20 aa and without oxidation, deamidation or potential missed cleavages were selected. Selected peptides were also confirmed with SRM Atlas database (www.srmatlas.org). To facilitate accurate relative quantification, synthetic heavy isotope-labeled peptides were obtained for all proteins. Survey unscheduled SRM assays with all possible y- and b-ion fragments were prepared for light and heavy peptides and monitored in testicular tissue or spermatozoa lysates on TSQ Quantiva^TM^. Intensity and interferences were assessed for each transition, and the three most intense transitions were selected for each heavy and light forms. Two separate multiplex SRM assays were finally developed for candidates identified in testicular tissues (20 heavy and light peptides for 9 candidates, and TEX101) and spermatozoa (20 heavy and light peptides for 9 candidates, and TEX101). All peptides were scheduled within 2-min intervals during a 30-min gradient (**Supplemental Tables S1 and S2**). TEX101-interacting proteins were verified in pools of independent testicular tissue and spermatozoa lysates (one biological replicate for each type of specimen). Three full process replicates were performed independently (from co-IP to trypsin digestion) for each specimen. Each process replicate was analysed in duplicate, and raw files were analyzed with Skyline software (v3.6.0.10493). The relative abundance of each endogenous peptide and corresponding protein was calculated according to the heavy-to-light ratio and the amount of the heavy peptides spiked in each sample. Non-specific mouse IgG antibody was used as a negative control. Proteins significantly co-enriched with TEX101 by *T1* antibody were confirmed by a two-sample *t*-test analysis of the mean fold change between the two groups (co-IP with *T1* and co-IP with non-specific mouse IgG). Cut-off values (fold change≥2, and p-value<0.01) were applied for the verification of the candidate TEX101-interacting partners in testicular tissue and spermatozoa.

### Protein digestion and SRM analysis for the verification of TEX101-interacting proteins

Co-IP of TEX101 complexes in testicular tissues and spermatozoa was performed, as described above. Prior to trypsin digestion, 500 fmoles of heavy isotope-labeled TEX101 and DPEP3 proteotypic peptides (AGTETAILATK*-JPTtag and SWSEEELQGVLR*-JPTtag) were added to all samples. Eight heavy isotope-labeled peptides for TEX101 interactome in testicular tissue, and eight heavy peptides for TEX101 interactome in spermatozoa, were pooled and diluted to a final concentration of 100 fmol/μL. Five μL of the heavy peptide pool were spiked to each sample after digestion. Initial testicular tissue and spermatozoa lysates (10 μg) were digested, to calculate the recovery of each protein after co-IP. Digests were desalted, and peptides were separated with a 30-min gradient and quantified by TSQ Quantiva^TM^ mass spectrometer. Peptides were loaded onto a 3 cm trap column (inner diameter 150 μm; New Objective, Woburn, MA, USA) packed in-house with 5 μm Pursuit C18 (Varian). An increasing concentration of Buffer B (0.1% formic acid in acetonitrile) was used to elute peptides from the trap column onto a resolving analytical 5 cm PicoTip emitter column (inner diameter 75 μm, 8 μm tip; New Objective) packed in-house with 3 μm Pursuit C18 (Varian). The SRM parameters were as follows: positive polarity, declustering and entrance potentials of 150 and 10 V, respectively; ion transfer tube temperature 300 °C; optimized collision energy values; scan time 20 ms; 0.4 and 0.7 Da full width at half maximum resolution settings for the first and third quadrupoles, respectively; and 1.5 mTorr argon pressure in the second quadrupole.

### Hybrid ELISA for the detection of TEX101-DPEP3 complex

Microtiter plates (96-well) were coated with anti-TEX101 antibodies (*T1* or *T2*; 500 ng per well). Following overnight incubation, plates were washed 3 times, and 100 μL of testicular tissue or spermatozoa lysates (prepared as previously described), or SP, were loaded on the plate. Two dilutions (10x and 4x for testicular tissue and spermatozoa lysate, and 100x and 10x for SP) in duplicates were used for each sample and each combination of antibodies. After 2 hours incubation with gentle shaking, plates were washed 3 times with PBS, and 100 μL of biotinylated anti-DPEP3 antibodies (*D1* or *D2*) were added to each well, and incubated for 1 hour. The plates were then washed with PBS and streptavidin-conjugated alkaline phosphatase was added for 15 min. After the final 6-times wash with PBS, 100 μL of DFP solution in substrate buffer were added and incubated for 10 min with gentle shaking. Finally, 100 μL of developing solution were added in each well for 1 min, and time-resolved fluorescence was measured. Reversed hybrid ELISAs were also performed simultaneously using anti-DPEP3 antibodies (*D1* and *D2*) for capture and biotinylated anti-TEX101 antibodies for the detection of TEX101-DPEP3 complexes. In addition, control experiments with non-specific mouse IgG (500 ng per well) for capture and all biotinylated anti-TEX101 or anti-DPEP3 antibodies for detection were performed simultaneously.

### Assessment of TEX101-DPEP3 complex disruption by anti-TEX101 and anti-DPEP3 monoclonal antibodies

In the first set of experiments, testicular tissue lysates were pre-incubated overnight with increasing concentrations (3.9 nM to 1000 nM) of *T2*, *T3*, *T4*, *D1*, and non-specific mouse IgG as a negative control, in duplicates. Hybrid immunoassay was performed to detect TEX101-DPEP3 complexes (**Supplemental Figure S1A**). Pre-incubation of testicular tissue lysate with *T4* and non-specific mouse IgG (15.5 nM to 5000 nM) and *D1* (15.6 nM to 1800 nM), followed by hybrid ELISA, was repeated in triplicates. In the second set of experiments, the format of the hybrid immunoassay was modified. Microtiter plates were coated with antibody *D2* (500 ng per well), and testicular tissue lysate was added to each well. Captured and purified complexes were incubated overnight with increasing concentration (0.01 nM to 1500 nM) of antibody *T4* and non-specific mouse IgG antibodies in triplicates. Detection antibody *T1* was added, and fluorescence was measured as described above. To determine the amount of total DPEP3 captured by *D2* in each well, regular DPEP3 ELISA was performed with *D2* as a capture and *D1* as detection antibodies. The One site – Fit logIC50 nonlinear regression model in GraphPad Prism (v5.03; Graphpad Software, San Diego, CA, USA) was used for curve fitting, and calculation of the half maximal effective concentration (EC50) for antibody *T4*.

### Assessment of O-sulfotyrosine modification in TEX101 protein

TEX101 protein was purified from testicular tissue lysate, spermatozoa lysate and SP using antibody *T1* or non-specific mouse IgG coupled to beads. Beads were washed and re-suspended in SDS-PAGE loading buffer (2x; BioRad, #1610737, Hercules, CA) with 5% β-mercaptoethanol, and heated at 95°C for 15 min. Original unpurified testicular tissue lysate, spermatozoa lysate and SP (10 μg total protein) were also included. Western blot analysis was performed with TEX101 (HPA041915, Sigma-Aldrich), and sulfotyrosine (sulfo-1C-A2) (Abcam, # ab136481) antibodies.

### Immunocapture-LC-MS/MS with mouse monoclonal anti-Sulfotyrosine antibody

Microtiter plates were coated with 500 ng/well of mouse monoclonal anti-sulfotyrosine antibody (sulfo-1C-A2) in 50 mM Tris buffer (pH 7.8). Antibody *T1* and non-specific mouse IgG were also used as positive and negative controls, respectively. Plates were washed, and then incubated with 10-fold diluted testicular tissue lysate (in 6% BSA), 10-fold diluted spermatozoa lysate or 100-fold diluted SP for 2 hours at RT. Plates were then washed with PBS (3 times) and 50 mM ABC (3 times), and samples were prepared for mass spectrometric analysis in Q Exactive™ Plus, as described above. Raw files were processed with MaxQuant software (version 1.5.2.8).

### Sample preparation and analysis by ImageStream flow cytometry

A fresh semen sample from a healthy fertile individual was collected and was allowed to liquefy at RT for 1 hour. The sample was centrifuged at 350 g for 5 min, and spermatozoa were then washed with PBS, and incubated with normal goat serum (NGS; 2%) for 25 min at RT. After blocking, *T1* and *D2* mouse mAbs (12.5 μg/mL) were added to the cell pellets, and samples were allowed to incubate for 2 hours at RT. After washing, Alexa Fluor 568^®^-conjugated secondary antibody (1 μg/mL) (goat anti-mouse IgG H&L Alexa Fluor 568^®^; ab175473, Abcam, Cambridge, MA) was incubated with cells for 1 hour at RT. Prior analysis, labeled spermatozoa were washed, and then incubated with the DNA binding dye Hoechst 33342 (Thermo Fischer Scientific, #H3570). Sperm cell pellet incubated only with Alexa Fluor 568^®^-conjugated secondary antibody was used as a negative control. Samples were analysed on an Amnis ImageStream Mark II, 5-laser two-camera Imaging Flow Cytometer (Amnis Corp., Seattle, WA). A bright-field (BF) area lower limit of 50 mm^2^ was used to eliminate debris and speed beads during acquisition, while detection channels included 1/9-for BF along with channels 4 and 7 for Alexa Fluor 568^®^ and Hoechst, respectively. Excitation was provided by the following laser lines and power settings: 405 nm (10mw), 561 nm (200mw) and 592 nm (200mw), while approximately 20,000 objects were captured for each sample using the low speed/high sensitivity settings at 60X magnification. Analysis was carried out using the IDEAS software supplied by Amnis.

### Data availability

Raw mass spectrometry data and MaxQuant output files were deposited to the ProteomeXchange Consortium via the PRIDE partner repository (www.ebi.ac.uk/pride/archive/login) with the dataset identifier PXD007515 and the following credentials: Username; Password: oj9BDrH1. SRM raw data were deposited to the Peptide Atlas repository with the dataset identifier PASS00990. URL: www.peptideatlas.org/PASS/PASS00990; Username: PASS00990; Password: UN5396gz; Full URL: ftp://PASS0090:UN5396gz@ftp.peptideatlas.org. Processed Skyline files can be downloaded at Panorama Public https://panoramaweb.org/TEX101proteincomplexes.url (, Password: C7xDe%75).

## RESULTS

### Production of TEX101 protein and mouse monoclonal antibodies recognizing its different epitopes

The mature form of human TEX101 protein was expressed in Expi293F cells. The peak of protein yield was acquired 72 hours after transfection. The expression and purity of TEX101 protein were evaluated by Coomassie staining SDS-PAGE and Western blot analysis using an anti-TEX101 rabbit polyclonal antibody (**Figure 1A**), and were also confirmed by mass spectrometry (**Supplemental Table S3**). Glycosylated forms of TEX101 were identified by mass spectrometry at ~29kDa (*band b*) and ~35 kDa (*band c*), while minor amounts of the non-glycosylated form were also detected at 20 kDa (*band a*) (**Figure 1A**). The purified recombinant TEX101 was quantified by an in-house TEX101 ELISA, as previously described (35).

**Figure 1.**
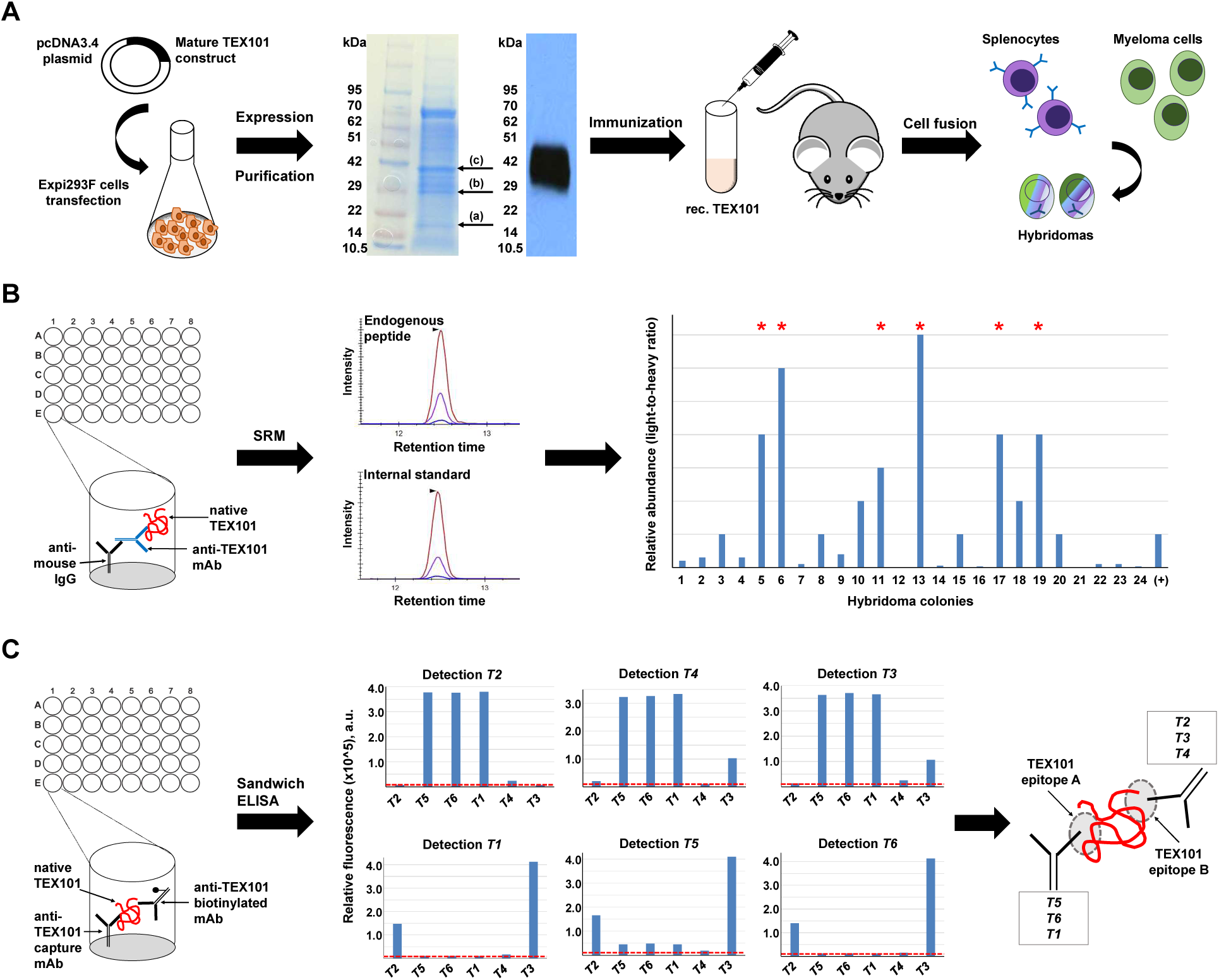
Production of mouse monoclonal antibodies against different epitopes of native TEX101 protein. **(A)** RhTEX101 protein was expressed by Expi293F cells. Western blot analysis with commercial rabbit polyclonal anti-TEX101 antibody (HPA041915), and SDS-PAGE followed by MS analysis confirmed the presence of purified TEX101 in the excised bands, marked by arrows (a-c). Mice were immunized with purified rhTEX101. **(B)** Immunocapture-SRM facilitated screening of hybridoma colonies and selection of mouse monoclonal antibodies against native TEX101 protein in the normal testicular tissue lysate. Lane (+) indicates anti-TEX101 mouse polyclonal antibody ab69522 used as a positive control. Asterisks mark the clones with enhanced binding to native TEX101. **(C)** Six mouse monoclonal anti-TEX101 antibodies were paired in sandwich immunoassays and revealed two groups of antibodies. Antibodies *T5*, *T6* and *T1* were directed against presumed Epitope A, while antibodies *T2*, *T4* and *T3* were directed against Epitope B. Dotted lines in red represent the background signal of sandwich immunoassays.

Mice immunization with the purified mature form of TEX101 generated 24 IgG-secreting hybridoma colonies. Hybridoma screening by immunocapture-selected reaction monitoring (SRM) revealed 12 hybridoma colonies producing antibodies that could capture native TEX101 from the testicular tissue lysate. Six out of 12 colonies produced antibodies with high affinity for native TEX101 protein (**Figure 1B**), and were subsequently expanded in serum-free media and purified using protein G columns. We showed previously that the commercial polyclonal antibody ab69522 could capture native TEX101 (28, 35). Immunocapture-SRM results revealed that our monoclonal anti-TEX101 antibodies possessed higher affinity for native TEX101 than ab69522 (**Figure 1B**). To investigate if in-house anti-TEX101 mAbs were directed against different TEX101 epitopes, we tested all possible combinations of capture and detection antibodies in a sandwich immunoassay. As a result, we identified two groups of antibodies, with each group targeting a different TEX101 epitope (**Figure 1C**). High affinity mAbs against multiple epitopes of the native endogenous TEX101 protein facilitated development of a coimmunoprecipitation-mass spectrometry (co-IP-MS) approach and thorough investigation of TEX101 physical interactome.

### Identification of the TEX101 physical interactome by co-IP-MS

To develop a stringent procedure for identification of TEX101 physical interactome, we optimized our sample preparation protocol and included mAbs against different epitopes of TEX101 and non-specific mouse IgGs as negative control. TEX101 interactomes were identified in testicular tissues, spermatozoa and SP.

Mild non-denaturing non-ionic (NP-40 and Triton X-100) and zwitterionic (CHAPS) detergents previously used for the solubilisation of membrane proteins in PPI studies (6, 7, 45) were tested for TEX101 isolation from testicular tissues. Following cryolysis, the highest recovery of TEX101 was achieved using CHAPS (1% w/v) for lysis and protein solubilization, as assessed by ELISA (**Supplemental Figure S2**). CHAPS sterol moiety could facilitate more efficient disruption of cholesterol-enriched lipid rafts and enhanced release of GPI-anchored complexes (46, 47). Antibodies were coupled to NHS-activated sepharose beads which previously revealed higher yields and lower non-specific binding in IP experiments (36). Since antibodies and TEX101-interacting proteins could compete for the same epitope, we selected two mAbs, *T1* and *T2*, generated against different TEX101 epitopes, as assessed by ELISA pairing (**Figure 1C**). Co-IP-MS experiments resulted in identification and relative quantification of several hundred proteins in testicular tissues, spermatozoa and SP. Proteins identified with false detection rate (FDR) of ≤1.0% were selected as putative TEX101-interacting proteins. Comparison of antibodies *T2* (160-fold enrichment of TEX101) and *T1* (616-fold enrichment of TEX101) in the testicular tissue lysate revealed the higher enrichment efficiency and higher yield of interacting proteins for *T1* antibody (**Figure 2A, Supplemental Figure S3**). Thus, *T1* was used for the enrichment of TEX101 complexes from spermatozoa and SP.

**Figure 2.**
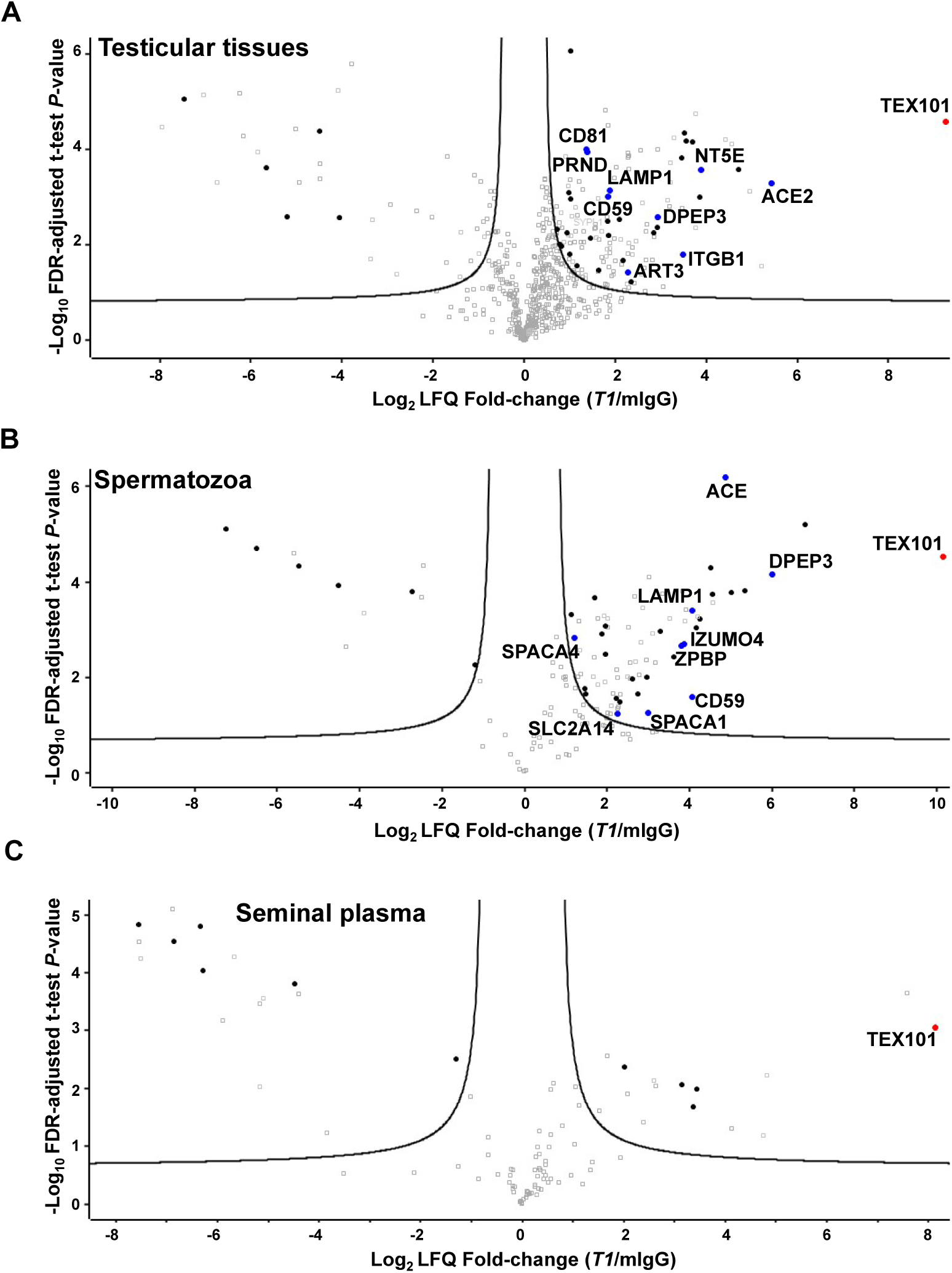
Identification of TEX101 protein interactome by co-IP-MS. Volcano plots revealed proteins co-enriched with TEX101 using *T1* antibody and testicular tissues with active spermatogenesis **(A)**, spermatozoa obtained from fertile individuals **(B)**, and pre-vasectomy seminal plasma **(C)**, as compared to the mouse IgG negative controls. Three biological replicates were used, and the hyperbolic curves indicate 1% FDR. Significantly enriched membrane-bound and secreted proteins are shown in black. Significantly enriched membrane-bound and secreted proteins expressed in testicular germ cells based on the Human Protein Atlas data (shown in blue) were subjected to verification in the independent sets of samples. Complete lists of proteins are presented in **Supplemental Tables S5, S6 and S7**.

Overall, 108 proteins were identified in testicular tissues with *T2* antibody at FDR≤1.0% and s0=0.27 (**Supplemental Table S4** and **Supplemental Figure S3**), and 135 proteins were identified with *T1* antibody at FDR≤1.0% and s0=0.29 (**Figure 2A, Supplemental Table S5**). Lists of candidates were filtered for secreted and membrane-bound proteins using HPA and NextProt databases (39 and 75 proteins for *T2* and *T1*, respectively). Examination of candidate expression in testicular germ cells narrowed down the number of proteins to 7 for *T2* antibody (**Supplemental Figure S3**) and 9 for *T1* (**Figure 2A and Table 1**). Seven proteins were found in common for *T2* and *T1* antibodies.

**Table 1.**
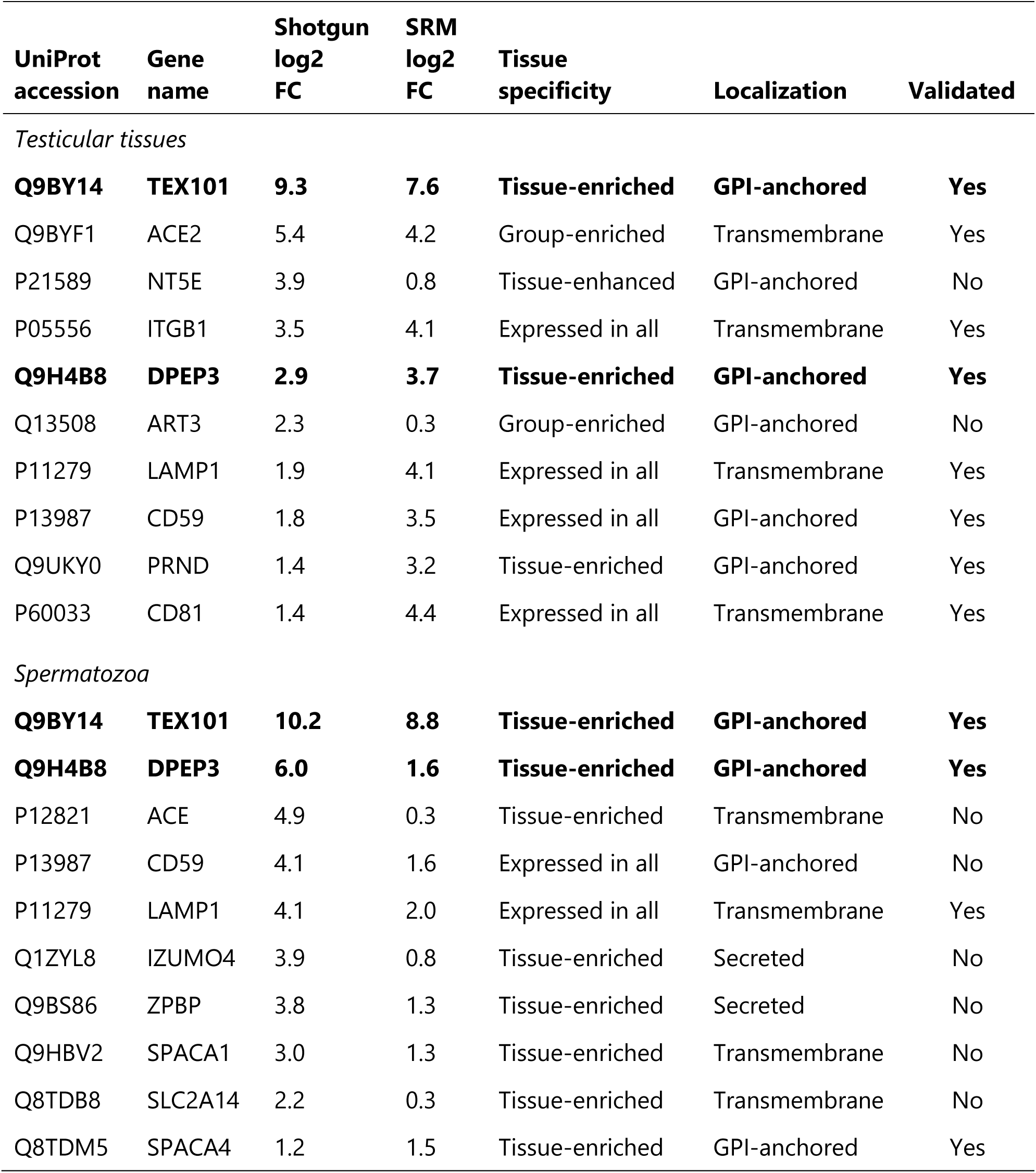
TEX101 interactome identified by co-IP-shotgun MS and validated by co-IP-SRM in the human testicular tissues and spermatozoa. FC, fold change.

Co-IP-MS in spermatozoa using *T1* antibody enriched TEX101 by 1,000 fold and identified 74 proteins at FDR≤1.0% and s0=0.60 (**Figure 2B, Supplemental Table S6**). Finally, 9 secreted and membrane-bound proteins were selected (**Table 1**). DPEP3, CD59 and LAMP1 proteins were common for tissues and spermatozoa lysates enriched with *T1* antibody. Comparison of candidates derived from tissues and spermatozoa suggested that *T2* antibody could share an epitope with TEX101-interacting proteins, and this competition could lead to the disruption of TEX101 complexes.

Co-IP-MS of soluble complexes in SP using *T1* antibody enriched TEX101 by 282-fold and identified 7 secreted and membrane-bound proteins at FDR≤1.0% and s0=0.58 (**Figure 2C, Supplemental Table S7**). Additional examination of these proteins revealed that 3 proteins were of epididymal origin, while 4 proteins were localized to intracellular membrane compartments. None of these 7 proteins were found in testicular tissues and spermatozoa. We thus concluded that TEX101 was present as a monomer in SP, which was in agreement with our previous findings (28).

### Verification of TEX101 interactome by co-IP-SRM

To verify TEX101 interactome, we used quantitative targeted mass spectrometry assays (48-51). Two multiplexed SRM assays in combination with co-IP were developed for monitoring the candidate proteins in testicular tissue and spermatozoa, respectively. We used an independent set of testicular tissue samples obtained from individuals with active spermatogenesis, and an independent set of spermatozoa samples from fertile individuals. We measured by SRM TEX101 protein, 9 candidate interacting proteins in testicular tissue and 9 candidate proteins in spermatozoa, before and after immunoprecipitation with *T1*. Overall, 7 out of the 9 candidate proteins were confirmed to be significantly (fold change≥2, and p-value<0.01) co-immunoprecipitated with TEX101 in testicular tissue (**Figure 3A and Table 1**), and 3 out of 9 candidates were confirmed in spermatozoa (**Figure 3B and Table 1**). Nearly 55% and 70% recovery of TEX101 protein with *T1* antibody in testicular tissue and spermatozoa was found after immunoprecipitation, respectively.

**Figure 3.**
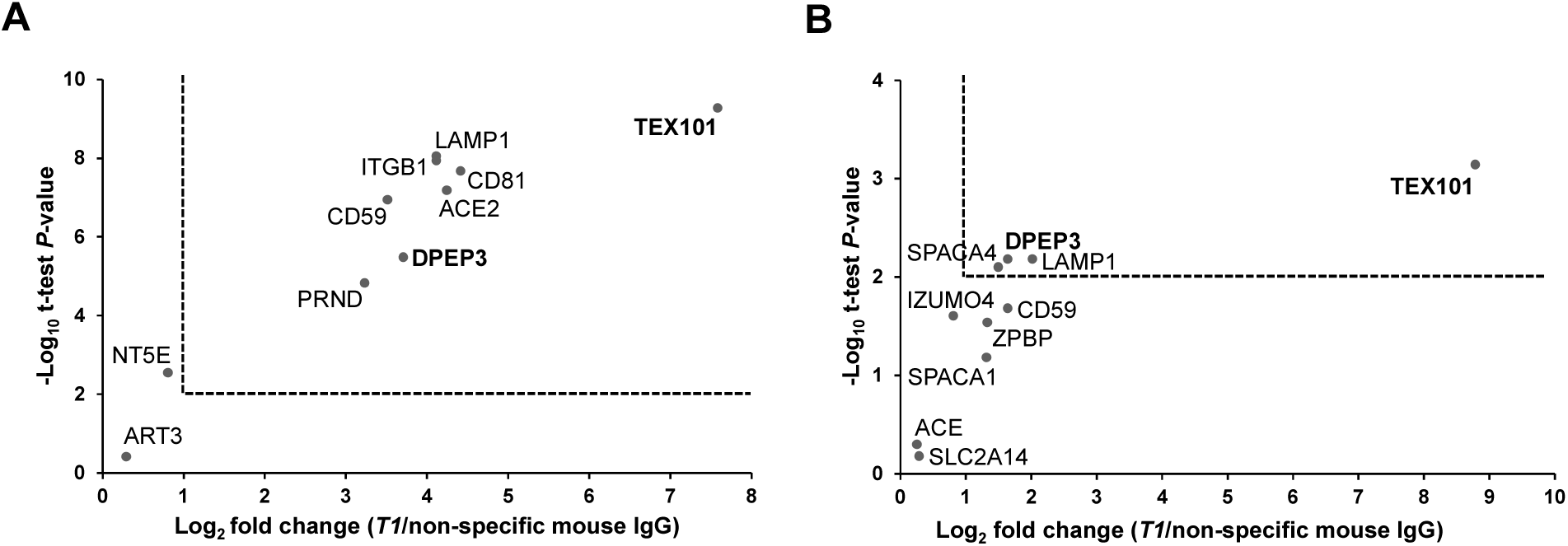
Verification of human TEX101 interactome by co-IP-SRM. Candidate proteins were measured in three independent testicular tissue **(A)** and spermatozoa **(B)** samples by multiplex SRM assays with heavy isotope-labelled peptide internal standards for the accurate relative quantification. Two-fold change and two-tailed *t*-test *P*-value = 0.01 were used as significance cut-offs (dotted lines).

### Production of DPEP3 protein and mouse monoclonal antibodies recognizing its different epitopes

Following examination of candidate proteins, we focused on dipeptidase 3 (DPEP3), a testis-specific GPI-anchored protein localized at the cell surface of testicular germ cells. DPEP3 expression pattern in human testicular germ cells was similar to TEX101, as assessed by HPA immunohistochemistry data (www.proteinatlas.org/ENSG00000141096-DPEP3/tissue). The mature form of human DPEP3 was expressed in Expi293F cells, and DPEP3 expression and purity were assessed by mass spectrometry (**Supplemental Table S8**), Coomassie staining, SDS-PAGE and Western blot analyses with anti-DPEP3 rabbit polyclonal antibody (**Supplemental Figure S4**). Purified rhDPEP3 was used as an immunogen for the production of mouse mAbs. Eight IgG-secreting clones were screened by IP-SRM for their ability to capture rhDPEP3 and native DPEP3 in SP, and two clones were selected (**Supplemental Figure S5A**), expanded in serum-free media and purified with protein G columns. Pairing these two anti-DPEP3 mAbs (*D1* and *D2*) in a sandwich format immunoassay showed that each antibody recognized a unique epitope of DPEP3 (**Supplemental Figure S5B**).

### Validation of TEX101-DPEP3 complex by a hybrid immunoassay

A TEX101-DPEP3 hybrid immunoassay was developed to confirm TEX101-DPEP3 complexes by independent orthogonal methods. Two anti-TEX101 (*T1* and *T2*) and two anti-DPEP3 (*D1* and *D2*) clones recognizing different epitopes were used as capture and detection antibodies, and vice versa. Hybrid ELISA confirmed TEX101-DPEP3 complexes in the testicular tissue and spermatozoa used for interactome discovery and validated the complex in independent testicular tissues and spermatozoa obtained from different patients. The hybrid ELISA also confirmed the absence of TEX101-DPEP3 complexes in SP. Based on signal intensity, the most efficient pair included *T1* and *D2* clones (**Figure 4A**; combinations 2 and 7). Combination of T2 and *D2* resulted in a lower signal (**Figure 4A**; combinations 4 and 8). Interestingly, combination of *T1* or *T2* with *D1* resulted in the loss of specific signal (**Figure 4A**; combinations 1 and 5, and 3 and 6). Hybrid ELISA with non-specific mouse IgG as capture antibody and all four biotinylated antibodies for detection revealed very low background fluorescence signal (**Supplemental Table S9**). Thus, hybrid ELISA confirmed the existence of TEX101-DPEP3 complexes in testicular tissues and spermatozoa, but not in SP.

**Figure 4.**
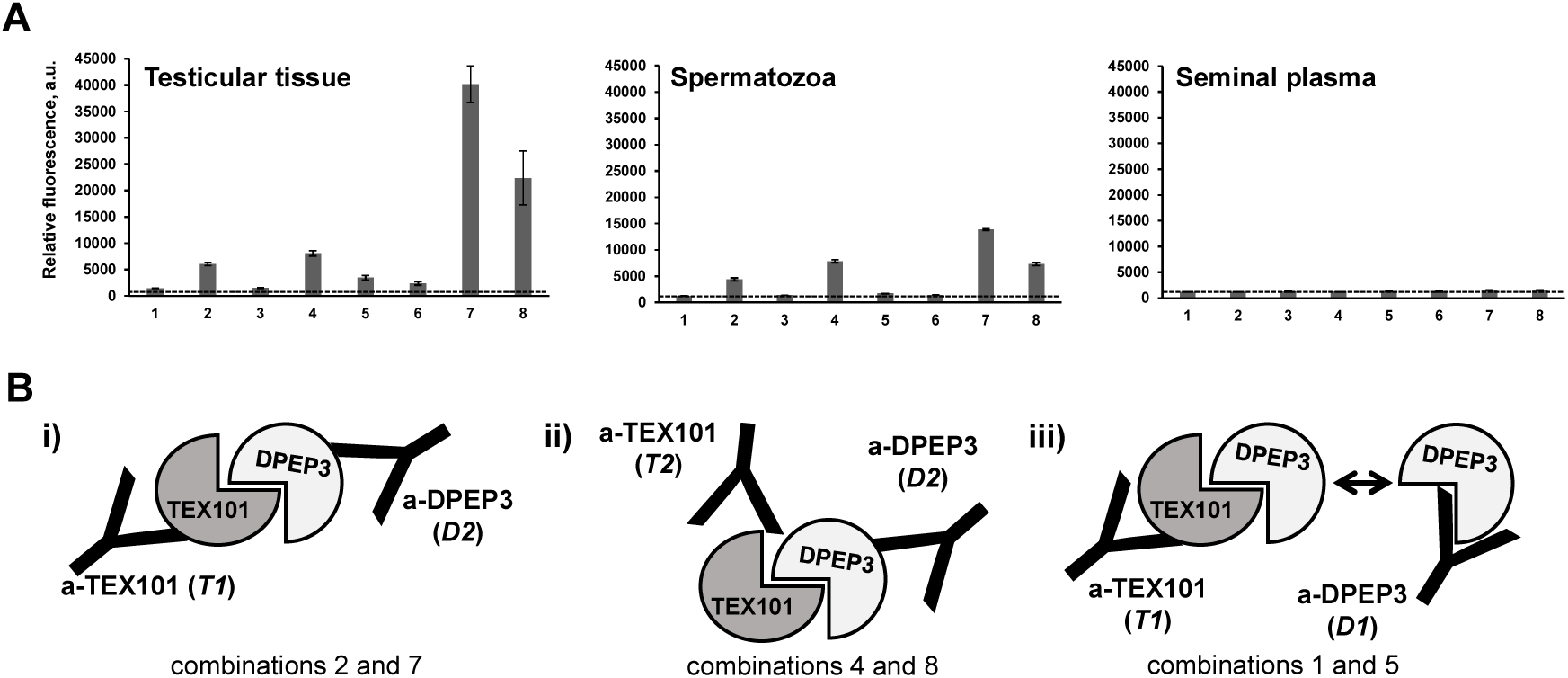
Validation of TEX101-DPEP3 complex by a hybrid immunoassay. **(A)** Relative abundance of TEX101-DPEP3 complex in an independent set of testicular tissue, spermatozoa and seminal plasma samples, as determined by TEX101-DPEP3 hybrid immunoassay. Various combinations of capture and detection monoclonal antibodies were used in a sandwich format: (1) *T1*–*D1*; (2) *T1*–*D2*; (3) *T2*–*D1*; (4) *T2*–*D2*; (5) *D1*–*T1*; (6) *D1*–*T2*; (7) *D2*–*T1*; (8) *D2*–*T2*. The mean fluorescence signal of the two replicates was calculated for 4-fold dilution of testicular tissue and spermatozoa lysates, and for 10-fold dilution of seminal plasma. Error bars represent the standard deviation of the two replicates, and dotted lines represent the background signal of TEX101-DPEP3 hybrid immunoassay obtained with non-specific mouse IgG for capture. **(B)** Schematic representation of monoclonal antibodies against different epitopes of TEX101 and DPEP3: **(i)** combination of *T1* and *D2* efficiently captured and detected TEX101-DPEP3 complex; **(ii)** combination of *T2* and *D2* was less efficient in detecting TEX101-DPEP3 complex, possibly due to partially accessible TEX101 epitope for *T2*; **(iii)** capture or detection by *D1* led to almost complete loss of fluorescent signal and suggested competition of *D1* for the same area of TEX101 binding.

### Assessment of tyrosine O-sulfation of TEX101 protein

To explain the absence of TEX101-DPEP3 complexes in SP, we hypothesized that TEX101-DPEP3 interaction on the surface of germ cells could be facilitated by a transient post-translational modification. Tyrosine O-sulfation has previously been identified as a post-translational modification which enhanced interaction of secreted and membrane-bound protein complexes (52). Protein sulfation was also crucial for sperm function and male fertility (5). For instance, tyrosylprotein sulfotransferase 2 (TPST2) knockout mice were infertile due to disruption and degradation of ADAM2-ADAM3 and ADAM2-ADAM6 complexes. It should be noted that these complexes were degraded in TEX101 knockout mice (7, 34). Thus, we investigated if human TEX101 was modified by tyrosine O-sulfation, and if such modification was crucial for stabilization of TEX101 complexes.

We assessed tyrosine O-sulfation by IP of TEX101 from testicular tissues, spermatozoa and SP followed by immunoblotting with anti-sulfotyrosine or anti-TEX101 antibodies (**Supplemental Figure S6**). As a result, TEX101 was enriched, but not detected by anti-sulfotyrosine antibody. In addition, IP-MS using anti-sulfotyrosine antibodies did not identify TEX101 in testicular tissues, spermatozoa or SP (**Supplemental Table S10**). We thus concluded that TEX101 was not modified by tyrosine O-sulfation. Further investigation of proteins modified by tyrosine O-sulfation may reveal the role of this post-translational modification in spermatogenesis and male fertility (53).

### Assessment of TEX101 and DPEP3 localization in human sperm cells

To confirm the localization of TEX101 and DPEP3 proteins in spermatozoa and immature sperm cells that were present in the semen, we used ImageStream flow cytometry and our monoclonal antibodies *T1* and *D2*. Two populations of cells, round germ cells (presumably haploid secondary spermatocytes) and mature spermatozoa, were identified and found positive for TEX101 and DPEP3 (**Figure 5**). Both TEX101 and DPEP3 proteins were localized to the cell surface of round germ cells (**Figure 5, A and B**). In mature spermatozoa, both TEX101 and DPEP3 were localized to the post-equatorial region of the sperm head (**Figure 5, D and E**). It has previously been shown that the post-equatorial region of sperm was involved in the sperm-egg interaction (54).

**Figure 5.**
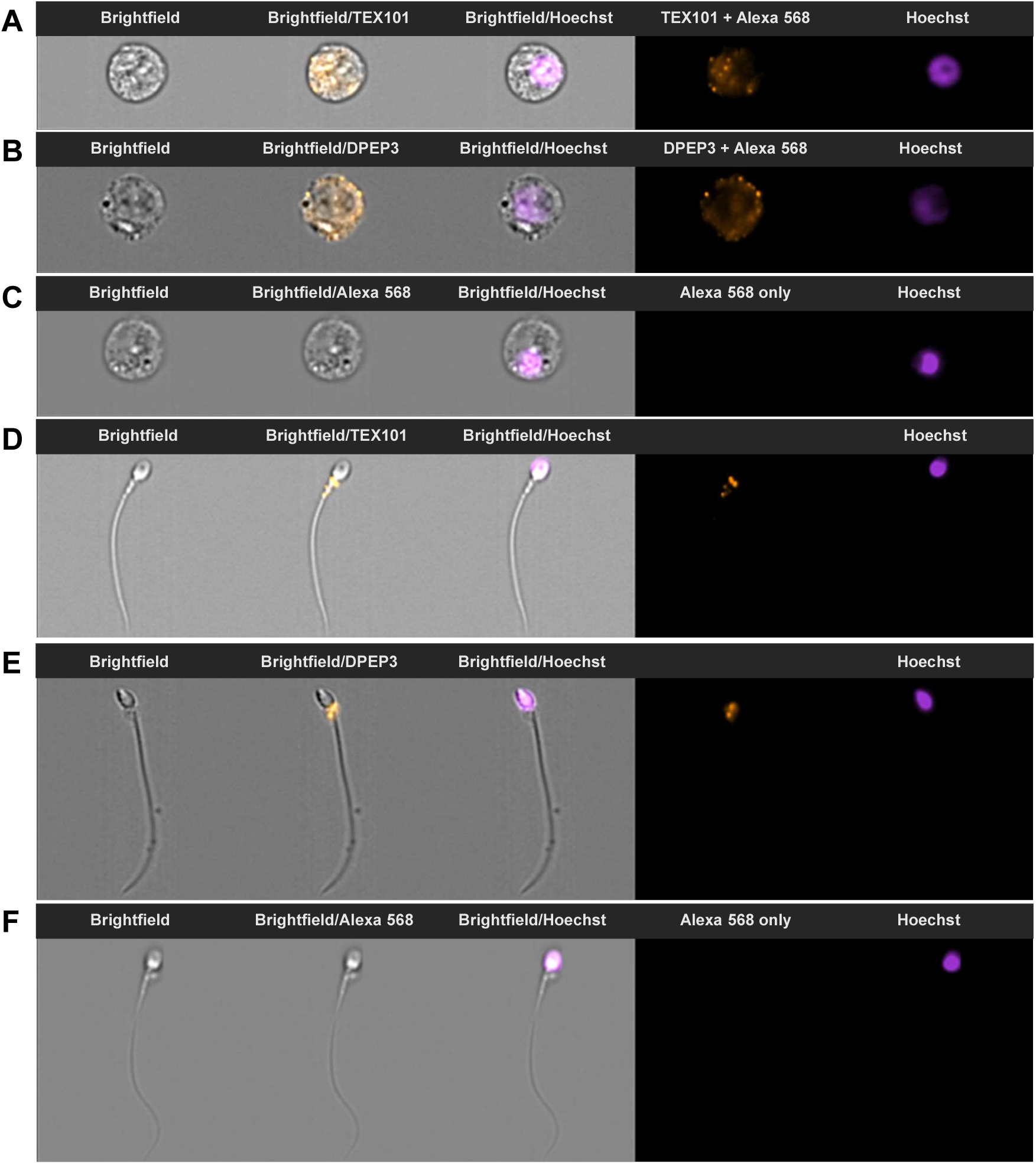
TEX101 and DPEP3 localization on germ cells and spermatozoa, as measured by imaging flow cytometry. During ImageStream analysis, single cells were separated into round germ cells (panels A-C) and mature spermatozoa (panels D-F) based on size and morphology. Counterstaining with Hoechst was used for nucleus visualization and discrimination between diploid and haploid cells. Panels A and B show secondary spermatocytes immunostained with anti-TEX101 *T1* and anti-DPEP3 *D2* antibodies, and detected with Alexa 568-conjugated goat anti-mouse secondary antibody. TEX101 and DPEP3 were localized to the cell surface. Panels D and E show spermatozoa immunostained with anti-TEX101 *T1* and anti-DPEP3 *D2* antibodies. TEX101 and DPEP3 staining was localized to the post-equatorial region of the sperm head. Panels C and F present round cells and spermatozoa incubated only with Alexa 568-conjugated goat anti-mouse antibody (negative control).

### Identification of antibody clones disrupting TEX101-DPEP3 complexes

TEX101-DPEP3 hybrid ELISA demonstrated that not all monoclonal antibodies against TEX101 and DPEP3 could capture TEX101-DPEP3 complex with equal efficiency. Combination of *D2* for capture and *T1* for detection was shown to be the most efficient antibody pair for detection of TEX101-DPEP3 complexes. Therefore, we assumed that both antibodies were directed against epitopes which were not involved in TEX101-DPEP3 interaction (**Figure 4; i**). Similarly, when pairing *T2* with *D2*, TEX101-DPEP3 complex was detectable, although fluorescence signal was approximately 50% lower compared to *D2* and *T1* combination. We thus assumed that antibody *T2* could not bind to TEX101 in the complex due to partially overlapping binding sites with DPEP3 protein (**Figure 4B (ii)**). Furthermore, hybrid ELISA signal was lost when *D1* anti-DPEP3 mAb was used to capture or to detect the fraction of DPEP3 in the protein complex. We hypothesized that antibody *D1* competed with the epitope occupied by TEX101 in the complex, and thus could capture only the free unbound DPEP3 (**Figure 4B (iii)**). Antibody *D1* was originally selected by its ability to bind recombinant DPEP3 or free soluble native DPEP3 present in SP. We thus assumed that antibody *D1* can be an inhibitory antibody and can potentially disrupt TEX101-DPEP3 complexes. We then re-analyzed a-DPEP3 and additional a-TEX101 clones and evaluated their disruptive efficiency. We first proceeded with overnight pre-incubation of increasing concentrations of selected clones with testicular tissue lysates followed by detection using *D2*/*T1* assay for a-TEX101 inhibitory antibodies, or *T1*/*D2* assay for a-DPEP3 inhibitory antibodies (**Supplemental Figure S1A**). Among all antibody clones, only *T4* and *D1* revealed dose-dependent decrease of fluorescence signal (**Supplemental Figure S1B**). We also then estimated that the amount of free unbound TEX101 and DPEP3 substantially exceeded the amount of TEX101-DPEP3 complexes in the testicular tissue lysate (~800 ng/mL free DPEP3 versus ~8 ng/mL TEX101-DPEP3 complexes). As a result, very high concentrations of antibodies were required to observe the decrease of fluorescent signal, and EC_50_ values were determined as 1080 nM [95%CI 454-2550] for *T4* and ~2000 nM for *D1* antibodies (**Supplemental Figure S1C**). As a result, we designed an assay, in which excess of free unbound TEX101 was washed away, while only TEX101-DPEP3 complexes were captured and then disrupted (**Figure 6A**).

**Figure 6.**
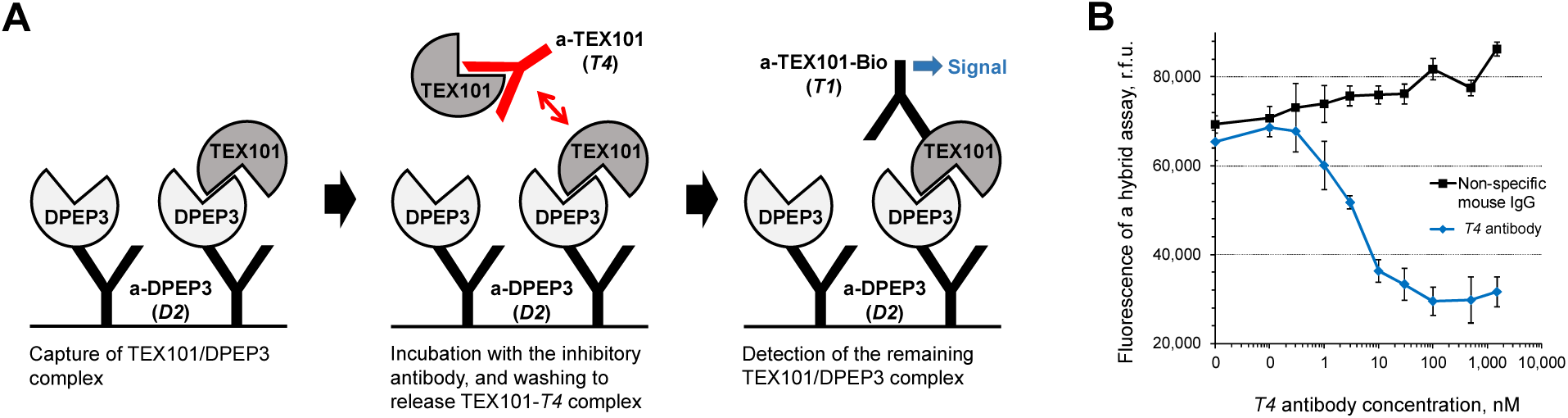
Hybrid immunoassay to screen for disruptors of TEX101-DPEP3 complex. **(A)** Schematic representation of hybrid immunoassay with anti-DPEP3 *D2* and anti-TEX101 *T1* antibodies. TEX101-DPEP3 complexes were captured and purified on the microtiter plates and then incubated with increasing concentrations of anti-TEX101 *T4* antibody. Relative abundance of the remaining TEX101-DPEP3 complexes were determined by fluorescence measurements. (B) Incubation of captured TEX101-DPEP3 complexes with anti-TEX101 *T4* antibody demonstrated a dose-dependent decrease of fluorescent signal. No decrease of signal was observed for a non-specific mouse IgG antibody. The EC50 value for anti-TEX101 *T4* was estimated at 3.4 nM [95%CI 2.4-4.9]. Error bars represent the standard deviation of the triplicates.

### Hybrid immunoassay to screen for candidate modulators of male fertility

With a new format of our hybrid assay, much lower concentration of *T4* and *D1* antibodies could disrupt TEX101-DPEP3 complexes. Clone *T4* had a much more profound effect, so we decided to focus on this clone. EC_50_ for T4 antibody was estimated at 3.4 nM [95%CI 2.4-4.9] (**Figure 6B**). Taking into account the amount of total DPEP3 captured from the testicular tissue lysate in each well (8.6 fmol in 100μL, or 0.086 nM) and assuming the affinity (K_d_) of antibody-protein (1:1) interaction as 1 nM, the affinity of TEX101-DPEP3 complex could be estimated as 40 pM [95%CI 26 - 57]. It should be noted, however, that we do not know the exact stoichiometry of *T4* antibody/TEX101 and DPEP3/TEX101 interactions.

We thus suggested that our hybrid ELISA with *D2* and *T1* antibodies could emerge as a simple but powerful platform to screen for molecules which disrupted TEX101-DPEP3 complexes.

## DISCUSSION

To investigate genes pertinent to spermatogenesis and fertilization, numerous knockout mouse models have been generated in past decades (8). Observed male infertility phenotypes were often associated with disrupted PPIs involved in sperm maturation, migration, zona pellucida binding and sperm-oocyte fusion (1-7). However, little knowledge on mouse testis-specific proteins has been translated into studies on human reproduction (13, 55), often due to the absence of human orthologs. For example, examination of testis-specific genes of the ADAM family revealed only six mouse genes with corresponding human orthologs (Adam2, Adam18, Adam21, Adam29, Adam30, Adam32), while twelve genes (Adam1a, Adam1b, Adam3, Adam4, Adam5, Adam6a, Adam6b, Adam24, Adam25, Adam26a, Adam26b, Adam34) did not have human orthologs or were non-coding pseudogenes in humans (56). Such difference between mouse and human genomes justified the studies on human testis-specific genes and proteins.

TEX101 is a prominent example of a highly testis-specific protein crucial for production of competent sperm and for fertilization (7, 34, 57-59). TEX101 function, as identified in mice, could be exerted through PPIs with numerous cell-surface testis-specific proteins. The most prominent mouse TEX101-interacting proteins Adam3, Adam5, Adam6a and Adam6b, however, are pseudogenes in humans, and ADAM4 is not present in the human genome. The roles of human TEX101 and its interactome thus remain unknown.

Previously, we reported on human TEX101 as a SP biomarker for the differential diagnosis of azoospermia (27, 60-62). We developed a first-of-a-kind TEX101 ELISA (35) and demonstrated its clinical utility in large cohorts of fertile, sub-fertile and infertile individuals (28). In the present study, we first focused on elucidation of the human TEX101 interactome in testicular tissues, spermatozoa and SP. We first optimized a co-IP-MS approach to ensure stringent identification of TEX101-interacting proteins. Choice of detergents was crucial since isolation of membrane GPI-anchored proteins and their complexes, often localized to cholesterol-enriched lipid rafts, is typically challenging due to their high hydrophobicity and resistance to detergents (63). Since our previous generation of monoclonal antibodies (35) could not efficiently enrich the native non-denaturated TEX101 from testicular tissues and SP, we produced second generation antibodies recognizing native TEX101. Our co-IP-MS approach identified and validated physical interactions of TEX101 in testicular tissues and in mature spermatozoa. Investigation of our candidates using the Contaminant Repository for Affinity Purification (64) revealed that none of our candidates were background contaminants. Interestingly, none of the testis-specific ADAM proteins (ADAM18, ADAM29 and ADAM32), the potential orthologs of mouse ADAM3-6 proteins, were found in the TEX101 interactome. This may suggest the transient nature of those interactions, or alternative mechanisms of spermatozoa maturation in humans. Identification of human TEX101 knockout or knockdown models, as well as more robust PPI studies involving protein cross-linking could be used to capture transient PPIs missed by co-IP-MS approaches (65).

Interestingly, no candidates emerged as TEX101-interacting proteins after co-IP-MS from SP (**Figure 2B**). Likewise, our TEX101-DPEP3 hybrid immunoassay validated the presence of TEX101-DPEP3 complexes in testicular tissues and spermatozoa, but not in SP (**Figure 4A**). These observations were in agreement with our previous size-exclusion chromatography data which revealed only the free soluble TEX101 in SP (28). Such differences between tissues, cells and SP could be the result of: (i) slightly alkaline pH 7.8 - 8.0 of SP weakening electrostatic interactions (66); (ii) loss of post-translational modifications or altered protein localization, and (iii) proteolytic degradation of TEX101-interacting proteins in SP. Here, we also demonstrated the absence of TEX101 tyrosine-O-sulfation, a recognized post-translational modification of extracellular PPIs (67) and interactions of testis-specific membrane proteins (5).

Literature review on TEX101-interacting proteins identified in this work revealed that DPEP3 has previously been shown to co-localize and form a physical complex with TEX101 on the surface of murine testicular germ cells (59). DPEP3 is a testis-specific membrane-bound protein of the dipeptidase family (68). Similarly to TEX101, DPEP3 is a GPI-anchored protein expressed by testicular germ cells. DPEP3 is shed into SP during sperm maturation (59). It was demonstrated that a fraction of murine DPEP3 in testicular tissues formed homodimers (59). Here, we demonstrated the presence of both DPEP3 monomers and homodimers in human testicular tissues and spermatozoa, while DPEP3 in SP was present as a homodimer (**Supplemental Figure S7**).

The enzymatic activity of DPEP3 was previously demonstrated *in vitro* (68), however, the molecular function of DPEP3 remains unknown. It could be suggested that TEX101 role as a cell membrane chaperone is exerted by TEX101-DPEP3 complexes, in which DPEP3 acts as a peptidase and cleaves pro-domains of ADAM proteins, while TEX101 modulates DPEP3 activity. Such hypothesis could be supported by formation of physical complexes and by localization to the GPI-protein enriched lipid rafts of the post-equatorial regions of spermatozoa. In our work, TEX101-DPEP3 complexes were detected by immuno-capture SRM and hybrid immunoassays in two and three different pools of tissue lysates, respectively, thus confirming the existence of this complex in different patients. It could be speculated that the signal observed in the hybrid immunoassays was not due to the existence of TEX101-DPEP3 complexes, but to the presence of an unknown interfering molecule. Such interfering molecule, however, should have at least three epitopes simultaneously recognized by monoclonal antibodies *D2*, *T1* and *T2*. Thus, the existence of such interference is unlikely. It is also unlikely that the dose-dependent decrease of hybrid immunoassay signal was due to competition between the disrupting clone *T4* and the detection clone *T1*, since these two clones were well-matched in a regular sandwich immunoassay (**Figure 1C**).

Suggested sub-nanomolar affinity of TEX101-DPEP3 makes it a relatively strong complex. With 179 complexes available in the Protein-Protein Interaction Affinity Database 2.0 (https://bmm.crick.ac.uk/~bmmadmin/Affinity), the affinity of complexes ranges from 24 fM to 635 μM, with the median affinity 13 nM. Further studies with purified TEX101 and DPEP3 proteins and known stoichiometries of interaction are needed to accurately measure the affinity of TEX101-DPEP3 complex and estimate the affinities of disrupting antibodies.

Finally, we suggest that the TEX101-DPEP3 hybrid immunoassay can emerge as a simple platform to screen for molecules which disrupt TEX101-DPEP3 complexes. We hypothesize that disruptors of TEX101-DPEP3 complexes could emerge as modulators of male fertility or male contraceptives. Even though we do not have data demonstrating that disruption of human TEX101-DPEP3 complexes *in vivo* leads to male sterility, such hypothesis could be supported by the following observations: (i) TEX101 and DPEP3 are proteins with very high testicular tissue and germ cell specificity and thus, should have unique roles in spermatogenesis and fertilization; (ii) TEX101 and DPEP3 proteins form a physical complex, as demonstrated in mice and human; (iii) both TEX101 and DPEP3 are GPI-anchored proteins localized to the lipid rafts and post-equatorial regions involved in the sperm-egg interaction; (iv) TEX101 knockout mice are sterile; (v) TEX101 has been shown to act as a pivotal chaperone for maturation and processing of ADAM proteins directly involved in sperm transit and sperm-egg interaction. Additional functional assays such as zona pellucida binding and hamster egg penetration assays may be required to validate the potential contraceptive effects of our a-TEX101- and a-DPEP3-disrupting antibodies.

With only few protein targets and molecular compounds proposed as modulators of male fertility and non-hormonal male contraceptives, the most promising compounds were either abandoned due to their side effects or are still under investigation in animal models (69-72). We believe that TEX101-DPEP3 complex may provide an alternative target to develop non-hormonal male contraceptives. Even though disruption of PPIs by small molecules or short peptides is challenging, it is not impossible (73). The ultimate male germ cell specificity of TEX101 and DPEP3 proteins would minimize potential side effects. With no oral non-hormonal male contraceptives available at the moment, the race for such molecules continues (74).

